# Benchmarking Agentic Large Language Models for Complex Protein-Set Functional Annotation

**DOI:** 10.64898/2026.04.18.719404

**Authors:** Xiaoyu Zhang

## Abstract

Large language model (LLM) agents are increasingly used to synthesize heterogeneous bioinformatics evidence, but their reliability for high-volume biological annotation remains poorly characterized. We evaluated three agent configurations on a controlled protein annotation task: Claude App with Claude Opus 4.7, Claude Code CLI with Claude Opus 4.7 and Claude Scientific Skills, and Codex App with GPT-5.4 and Claude Scientific Skills. Each configuration was run three times on the same verbatim prompt, the same 73 selected orthogroup FASTA files (1,705 protein sequences), and the same local evidence: Swiss-Prot BLASTP output, Pfam/HMMER domain hits, DeepTMHMM topology predictions, and SignalP secretion predictions. We audited the nine outputs for coverage, biological correctness, missing evidence, hallucinated or over-specific annotations, and within-method consistency, then merged the best-supported evidence into a final orthogroup annotation table. All nine runs covered all 73 orthogroups, indicating that the agents could retrieve and organize the complete input set. However, normalized calcification-relevance calls were only moderately reproducible: within-method exact tier agreement ranged from 0.397 to 0.685 for Claude App (mean 0.562), 0.342 to 0.740 for Claude Code (mean 0.516), and 0.411 to 0.630 for Codex App (mean 0.539), and the per-run number of high-confidence calls varied from 0 to 12 across the nine runs. The final curated table retained 3 high-confidence, 9 moderate, 18 watchlist, and 43 low-relevance orthogroups. The most robust direct candidates were sulfatase (OG0017138) and sulfotransferase (OG0020703) families and an FG-GAP/integrin-like surface protein family (OG0018986), whereas common error modes included elevating pentapeptide-repeat orthogroups on motif evidence alone, treating weakly secreted housekeeping enzymes as matrix proteins, and taking low-complexity BLAST labels at face value. Skill-enabled agents improved file handling, evidence traceability, and reproducibility of computational checking, but they did not eliminate biological overinterpretation. These results support a best-practice workflow in which LLM agents draft annotations only after deterministic evidence tables are generated, with explicit scoring rules, provenance columns, run-to-run replication, and expert review of high-impact claims.

## 1 Introduction

Protein function annotation is a persistent bottleneck in comparative genomics. Sequence databases now contain hundreds of millions of protein records, and even curated resources combine expert review, automated rule transfer, domain signatures, and machine-learning predictions to keep pace with new genomes [1]. Homology searches such as BLAST remain a practical first pass [2], while profile-HMM domain searches against Pfam add family-level evidence that is often more sensitive for remote homologs [3, 4]. Secretion and membrane-topology predictors such as SignalP 6.0 and DeepTMHMM provide orthogonal context about whether a protein could act in extracellular matrices, vesicles, or membranes [5, 6]. Combining these evidence streams is routine, but the final biological interpretation is still difficult because many protein families are multidomain, repetitive, lineage-specific, or only indirectly connected to the process of interest. Community-wide evaluations of automated function prediction have also shown that even the best classifiers fall well short of perfect agreement with expert-curated annotations, with reliability that varies systematically across functional categories and taxonomic groups [7].

The task considered here is especially challenging: annotating selected orthogroups for possible relevance to calcification. Coccolithophore calcification involves CaCO_3_ scale formation, endomembrane trafficking, ion and proton handling, and organic matrix components [8, 9]. Coccolith-associated polysaccharides can contain acidic and sulfated residues and influence calcite crystal morphology [10], and recent proteomic surveys of isolated coccoliths have begun to catalog matrix-associated proteins, including those carrying pentapeptide-repeat motifs that may help direct calcite formation [11]. Therefore, sulfation, glycosylation, secreted matrix proteins, adhesion proteins, transporters, and vesicle-associated factors can all be biologically plausible. At the same time, the same vocabulary creates a high risk of false positives: a generic secreted enzyme, a single weak collagen-like BLAST hit, or a calcium-binding domain in an unrelated intracellular protein should not automatically be called a calcification factor.

Agentic LLM systems appear attractive for such annotation because they can read many files, summarize evidence, write reports, and in coding environments run scripts to check their own outputs. Claude Code is described by Anthropic as an agentic coding tool that can read a codebase, edit files, run commands, and integrate with development tools [12]. OpenAI describes Codex as a coding agent that can read, edit, and run code, including background and parallel cloud tasks [13]. Scientific skill libraries add domain-specific instructions, examples, and workflow scaffolds; the K-Dense Scientific Agent Skills repository currently describes a collection of ready-to-use scientific and research skills for agent systems [14]. These capabilities may improve the mechanics of scientific workflows, but they do not by themselves guarantee biological correctness.

Here we use a controlled, repeated-run comparison to evaluate three agent configurations as they were used on the same orthogroup annotation prompt. The goal is not to rank proprietary foundation models in general. Instead, we ask a practical question relevant to bioinformatics groups: when agents are asked to retrieve, integrate, and interpret a large set of complex protein annotations, where do they help, where do they fail, how consistent are repeated runs, and how should their outputs be merged into a defensible final annotation table?

## 2 Results

### 2.1 All agents completed the retrieval task, but interpretation varied substantially

Each of the nine runs returned annotations for all 73 orthogroups, so no method failed at the basic retrieval and coverage layer. The shared input comprised 1,705 protein sequences, 73 BLAST tables, 73 Pfam/HMMER tables, seven DeepTMHMM topology batches, and two SignalP batches. The downstream differences therefore reflected how the agents interpreted evidence rather than whether they found the files.

The final expert-audited merge classified 3 orthogroups as high relevance, 9 as moderate relevance, 18 as watchlist candidates, and 43 as low relevance (Figure 1). The high-confidence set was intentionally conservative. It included OG0017138, an arylsulfatase/sulfatase family; OG0020703, a heparan-sulfate or sulfoglycan sulfotransferase-like enzyme; and OG0018986, an FG-GAP/integrin-alpha-like cell-surface receptor family with secretion or membrane-topology support. Moderate candidates were retained when evidence supported an indirect or plausible matrix, polysaccharide, surface, or remodeling role but lacked enough family-wide specificity for a high-confidence call.

**Figure 1:**
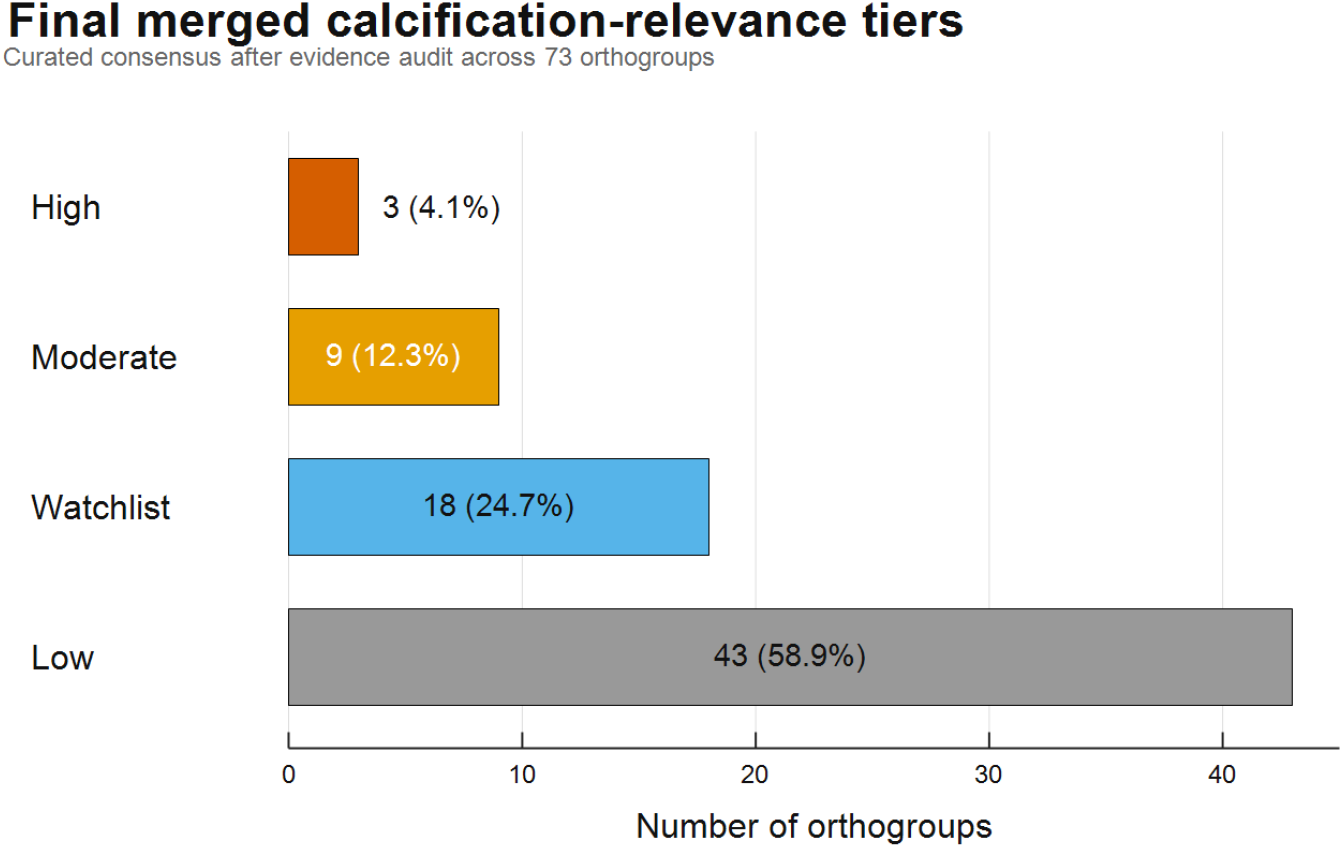
Final curated calcification-relevance distribution after the controlled benchmark workflow. The shared input consisted of 73 orthogroup FASTA files with Swiss-Prot BLASTP, Pfam/HMMER, DeepTMHMM, and SignalP evidence. Three independent runs were performed for each agent configuration: Claude App Opus 4.7, Claude Code Opus 4.7 with scientific skills, and Codex App GPT-5.4 with scientific skills. Model outputs were normalized to common relevance tiers, checked against recomputed evidence summaries, audited for overcalls and missing evidence, and merged into the final annotation table.

### 2.2 Run-level calibration differed within and between methods

Although coverage was complete, calibration differed strongly across runs (Figure 2; Table 1). Claude App run 2 was highly conservative, assigning 67 of 73 orthogroups to a low or background tier and only one high call. Claude App runs 1 and 3 were less conservative, with 11 and 8 high calls, respectively. Claude Code with scientific skills produced fewer high calls overall (1, 3, and 2), but shifted substantially between low and watchlist labels across runs. Codex App with scientific skills showed the widest high-call range, from no high calls in run 2 to 12 high calls in run 3.

**Table 1:** Run-level relevance calibration after normalization of each output to common evidence tiers.

| Run | High | Moderate | Watchlist | Low | Mean score | Main calibration pattern |
| --- | --- | --- | --- | --- | --- | --- |
| Claude_app_run1 | 11 | 15 | 20 | 27 | 1.71 | Broadly permissive; many secreted or membrane families promoted. |
| Claude_app_run2 | 1 | 5 | 0 | 67 | 0.68 | Most conservative output; few indirect candidates retained. |
| Claude_app_run3 | 8 | 10 | 10 | 45 | 1.29 | Intermediate, but still promoted several weak matrix candidates. |
| Claude_code_run1 | 1 | 10 | 0 | 62 | 0.82 | Conservative and similar to **Codex_app_run1**. |
| Claude_code_run2 | 3 | 15 | 22 | 33 | 1.36 | Better separation of watchlist from low, but still some overcalls. |
| Claude_code_run3 | 2 | 16 | 34 | 21 | 1.50 | Most watchlist-heavy Claude Code output. |
| Codex_app_run1 | 1 | 10 | 0 | 62 | 0.82 | Conservative baseline-like output. |
| Codex_app_run2 | 0 | 34 | 0 | 39 | 1.43 | Broad plausible-indirect labeling with no high tier. |
| Codex_app_run3 | 12 | 34 | 0 | 27 | 2.01 | Most permissive run; many indirect candidates promoted. |

**Figure 2:**
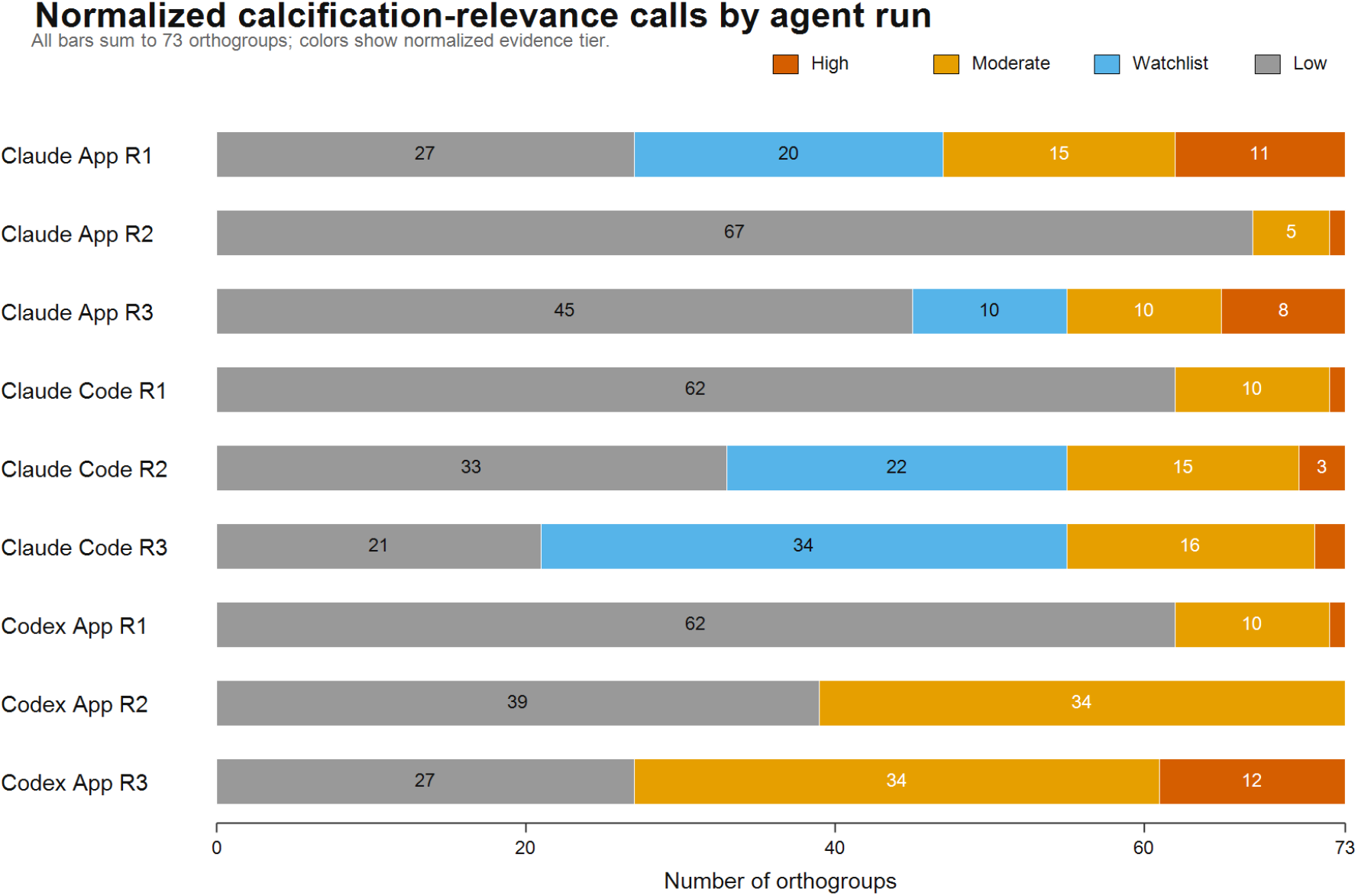
Normalized calcification-relevance distributions for the nine agent runs. Each horizontal bar sums to the 73 shared orthogroups, showing that all runs had complete coverage but differed substantially in calibration.

Two additional calibration patterns stood out. First, Codex App never emitted a watchlisttier label in any of its three runs, instead splitting every orthogroup among high, moderate, and low tiers; this is a vocabulary choice rather than a coverage failure, but it forced the normalization step to re-bucket moderate Codex calls that functionally behaved like watchlist candidates, and it prevented Codex from marking orphan-secreted or orphan-membrane families as explicitly provisional. Second, Claude Code run 1 and Codex App run 1 produced identical normalized distributions (high=1, moderate=10, low=62), despite coming from different model families and different agent interfaces; this apparent convergence on a conservative baseline under stochastic decoding suggests that when the prompt is interpreted literally, both agent families default to a similar “only call what is directly evidenced” pattern.

Within-method exact agreement on normalized relevance labels was modest (Figure 3; Table 2). The best agreement was between Claude Code runs 2 and 3 (54/73 orthogroups; 0.740), while the lowest was between Claude Code runs 1 and 3 (25/73; 0.342). Mean within-method agreement was in the same range for all three configurations (0.516–0.562), so no configuration was dramatically more reproducible than the others at the tier-label level. These results argue against relying on a single stochastic agent run for final biological claims, even when the input files and prompt are identical.

**Table 2:** Within-method exact agreement between repeated runs after tier normalization.

| Method | Run-pair agreements | Mean agreement | Range |
| --- | --- | --- | --- |
| Claude App Opus 4.7 | 0.397, 0.603, 0.685 | 0.562 | 0.397–0.685 |
| Claude Code Opus 4.7 + skills | 0.466, 0.342, 0.740 | 0.516 | 0.342–0.740 |
| Codex App GPT-5.4 + skills | 0.630, 0.411, 0.575 | 0.539 | 0.411–0.630 |

**Figure 3:**
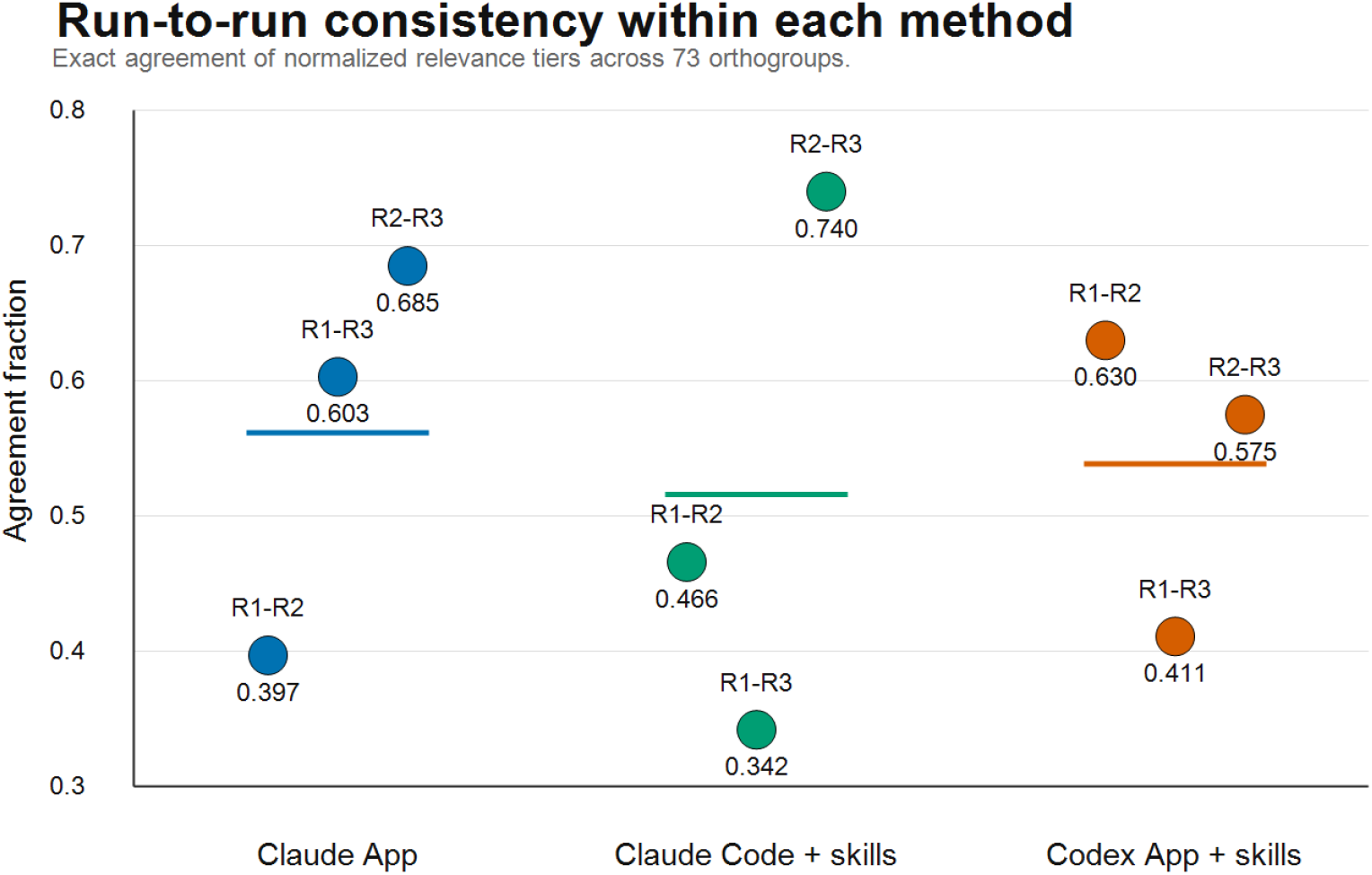
Pairwise run-to-run consistency within each method. Dots show exact normalized-tier agreement for each pair of repeated runs; horizontal lines show the method mean.

### 2.3 Biologically robust candidates shared direct matrix-chemistry or surface evidence

The final high-confidence candidates were those for which the evidence connected directly to known features of calcification biology rather than only to generic secretion or membrane localization (Table 3). OG0017138 had strong family-wide sulfatase support, with 14 of 16 sequences carrying a Sulfatase Pfam hit (best *E*-value 4.6 × 10^−48^) and all 16 sequences mapping to arylsulfatase BLAST hits. OG0020703 combined a coherent though more sparsely covered sulfotransferase signal (1 of 11 sequences with a Sulfotransfer_1/Sulfotransfer_3 Pfam hit and 2 of 11 with heparan-sulfate-6-*O*-sulfotransferase BLAST hits) with family-wide secretion support (4/11 SignalP-positive; no predicted transmembrane regions) in an orthogroup size small enough that a family-level sulfotransferase assignment is reasonable. These functions are biologically coherent because coccolith-associated polysaccharides include acidic and sulfated components that can influence calcium carbonate crystallization [10]. OG0018986 was retained as high because it combined family-wide FG-GAP repeats (12 of 13 sequences with FG-GAP and FG-GAP_3 Pfam hits) with extracellular or membrane topology (7/13 SignalP-positive, 10/13 with predicted transmembrane or SP+TM topology). This is not a direct enzyme in carbonate chemistry, but it is one of the strongest non-enzymatic surface or adhesion candidates in the dataset.

**Table 3:** Final high and moderate calcification-relevance candidates after evidence audit.

| Tier | Count | Orthogroups |  | Evidence basis |
| --- | --- | --- | --- | --- |
| High | 3 | OG0017138;<br>OG0020703 | OG0018986; | Direct sulfatase or sulfotransferase chemistry tied to sulfated matrix glycans, plus one FG-GAP/integrin-like surface family with strong topology and domain support. |
| Moderate | 9 | OG0009246;<br>OG0009816;<br>OG0017305;<br>OG0023594;<br>OG0025059 | OG0009301;<br>OG0011061;<br>OG0021347;<br>OG0024846; | Secreted adhesive or orphan proteins, glycosyltransferase-like enzymes, sulfotransferase with limited coverage, and secreted protease or proline/cysteine-rich matrix candidates. |
| Watchlist | 18 | See final TSV |  | Orphan secreted or membrane families, membrane organizers, signaling proteins, or weakly supported matrix-like families requiring follow-up evidence. |
| Low | 43 | See final TSV |  | Housekeeping, intracellular, weakly annotated, or unsupported families without calcification-specific evidence. |

The moderate tier was deliberately heterogeneous. For example, OG0009816, OG0011061, OG0023594, and OG0024846 were glycosyltransferase-like candidates with plausible links to matrix or cell-wall polysaccharide biosynthesis, but their evidence was too indirect or incomplete for a high call. OG0025059 was a strong novel secreted orphan candidate, with five of seven SignalP-positive sequences and no transmembrane prediction, but it lacked a functional domain. Such proteins are valuable experimental targets but should be annotated as candidates rather than known calcification proteins.

The evidence heatmap in Figure 4 shows why the final high and moderate candidates could not be treated as a single homogeneous class. Some orthogroups had strong family-wide BLAST and Pfam support, while others were retained primarily because of secretion or membrane topology. The heatmap also highlights cases with high model consensus but limited domain coverage, which were the cases most likely to require manual downgrading.

**Figure 4:**
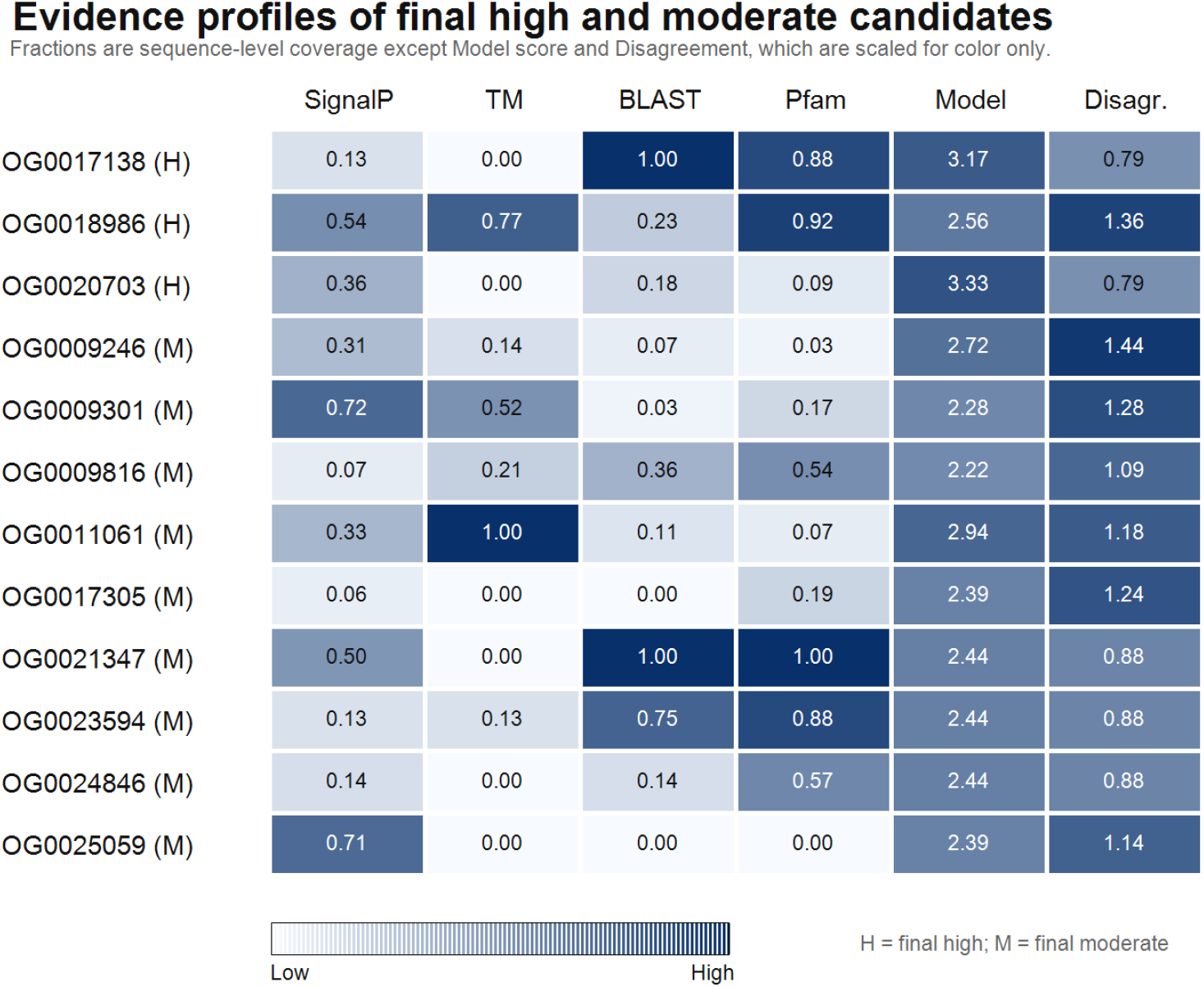
Evidence profiles for final high and moderate candidates. SignalP, TM, BLAST, and Pfam columns show sequence-level fractions. Model is the mean normalized model relevance score, and disagreement is the between-run standard deviation; these two columns are numerically labeled with their raw values but color-scaled for display.

### 2.4 Common hallucination and overclaim modes were biologically interpretable

The most important error mode was not invented file content but over-specific biological inference from real but weak evidence. Pentapeptide-repeat orthogroups (OG0001976, OG0010867, and OG0022524) were elevated by some runs on the basis of pentapeptide-repeat signal alone, with mostly globular topology and no direct calcium- or carbonate-binding marker. At the level of protein-family biochemistry, pentapeptide-repeat proteins are characterized by a right-handed quadrilateral *β*-helical (Rfr) fold and have no intrinsic calcium- or carbonate-binding activity [15], so agents that jumped directly from “pentapeptide repeats” to a calcification function were over-reaching the local evidence. However, a recent proteomic survey of *Emiliania huxleyi* coccolith matrix proteins identified pentapeptide-repeat motifs as a recurring feature of matrix-associated proteins and suggested, by analogy to the aspein pearl-oyster matrix protein, that these repeats may contribute to directing calcite formation during coccolithogenesis [11]. We therefore treat the pentapeptide-repeat orthogroups as defensible watchlist candidates rather than clear positives: assigning them directly to a mechanistic calcification role remains unsupported by the per-orthogroup evidence in this benchmark, but dismissing them outright also ignores that matrix-proteomic surveys place the motif inside the coccolith matrix. Several secreted housekeeping enzymes were similarly overranked when secretion was treated as sufficient evidence for matrix function. Low-complexity or repetitive BLAST hits also misled some outputs: the DNA-directed RNA polymerase II subunit RPB1 hit for OG0009301, Piccolo-like hits for OG0011061 or OG0015153, and a collagen call for OG0016203 based on one member of an 18-sequence family all required downgrading.

These mistakes mirror known issues in automated protein annotation. Public databases and computational transfer pipelines can accumulate overprediction and misannotation when specific functions are assigned beyond the support provided by family-level evidence [16]. LLM agents add an additional layer: they can produce fluent rationales that make weak annotation transfer sound mechanistically complete. For this dataset, the safer interpretation rule was to annotate at the specificity level actually supported by BLAST, Pfam, topology, secretion, and family-wide coverage.

### 2.5 Scientific skills improved workflow mechanics more than biological calibration

This benchmark cannot isolate skills as an independent causal variable because the skill-enabled conditions also used different interfaces from Claude App, and Codex used a different model family. Nevertheless, the qualitative comparison is informative. The skill-enabled coding-agent runs were better suited to deterministic checks: they could inspect folders, aggregate evidence, write scripts, and produce reproducible TSV outputs. That advantage was important for coverage checking and provenance, and it made it easier to diagnose disagreements after the fact.

However, the presence of scientific skills did not guarantee conservative biological judgment. Claude Code produced both conservative and watchlist-heavy outputs, while Codex ranged from conservative to highly permissive. This suggests that skills help agents use tools and domain workflows, but final biological calibration still depends on explicit prompt constraints, scoring rules, and review. In other words, skills improved the annotation process, but not enough to replace an evidence audit.

## 3 Discussion

The primary lesson from this benchmark is that agentic LLMs are useful assistants for protein-set annotation, but they should be treated as evidence organizers and hypothesis generators, not autonomous curators. All three agent configurations found and processed the complete orthogroup set. This is a meaningful practical success for large annotation folders, where failures often arise from missed files, inconsistent filenames, or incomplete output tables. The agents also surfaced plausible biology, especially sulfation, glycosylation, secreted orphan proteins, and membrane candidates.

The main weakness was calibration. The same prompt and same evidence produced runs that ranged from one high-confidence candidate to twelve. The final manual merge retained only three high-confidence orthogroups, indicating that several runs overestimated calcification relevance. The most permissive outputs were not useless; they often identified good watchlist candidates. Their problem was that they blurred the distinction between direct evidence, indirect pathway plausibility, and speculative follow-up.

For similar bioinformatics tasks, we recommend a staged workflow. First, generate deterministic evidence tables before asking an LLM for interpretation. Each orthogroup should have sequence counts, species counts, BLAST hit coverage, Pfam hit coverage, best E-values, SignalP fractions, DeepTMHMM topology summaries, and any low-complexity or single-member caveats. Second, require agents to assign both a function label and an evidence tier, with explicit definitions for high, moderate, watchlist, and low. Third, run the same prompt multiple times or across agents, then inspect disagreements rather than averaging them away. Fourth, include a negative-control mindset: housekeeping enzymes, single-hit collagen labels, repetitive proteins, and generic secretion should be downgraded unless supported by family-wide domains or process-specific biology. Finally, reserve final labels such as “calcification protein” for experimentally supported genes or very strong mechanistic evidence; use “candidate”, “watchlist”, or “indirectly relevant” for the more common cases.

The comparison also suggests that skill libraries are most valuable when they push agents toward reproducible computation. Skills should include templates for evidence schemas, scripts for re-aggregating annotation files, checks for missing orthogroups, and prompts that force conservative interpretation of weak homology. They should not only provide domain vocabulary, because vocabulary without calibration can increase overconfident mechanistic narratives.

### 3.1 Limitations

Several limitations of this benchmark should be kept in mind when generalizing from the results. First, the three agent configurations differ in more than one axis at once: Claude App uses a chat-style interface without a local code-execution environment, while Claude Code and Codex App run in coding-agent interfaces with local file and shell access, and only two of the three conditions use the scientific skill library. We therefore cannot cleanly attribute observed differences to model family, interface, or skill library as independent factors. Second, we ran each configuration only three times, which is sufficient to show that single-run biological conclusions are unstable but too small to support quantitative inference on reproducibility differences between configurations; larger replicate counts would be required to claim, for example, that Claude App is reliably more or less consistent than Codex App. Third, the benchmark target is calcification relevance in one taxonomic context, and the error modes observed here (elevating pentapeptide-repeat orthogroups from local evidence alone, treating secretion of housekeeping enzymes as matrix function, and taking low-complexity BLAST labels at face value) may be more or less prominent in other biological tasks. Fourth, the curated merge is itself an expert judgment rather than a held-out ground truth, so “overcalling” and “undercalling” should be read relative to a defensible conservative standard, not as errors against a gold label. Fifth, the relevance categories we report are ordinal but coarse; a finer-grained scoring rubric, or quantitative evidence-weight features, could reduce inter-run variability that comes purely from threshold drift between runs.

## 4 Methods

### 4.1 Input data and agent configurations

The benchmark used the folder `Orthogroups.calcifying_loose_fastas`, which contained 73 selected orthogroup FASTA files spanning 1,705 protein sequences from a comparative coccol-ithophore proteome. Each method received the same verbatim prompt asking it to annotate the biological function of each orthogroup, assess relevance to calcification, and use the local BLASTP, Pfam/HMMER, DeepTMHMM, and SignalP outputs in the same input directory; agents were also permitted to generate additional supporting evidence if they judged it necessary. The three evaluated configurations were: Claude App with Claude Opus 4.7 [17] at the highest available reasoning setting; Claude Code CLI with Claude Opus 4.7 at the highest available reasoning setting and the Claude Scientific Skills library; and Codex App with GPT-5.4 at the highest available reasoning setting and the same scientific skill library. Each configuration was run three independent times with no inter-run state, conversation history, or shared scratchpad.

### 4.2 Evidence audit and normalization

After the nine outputs were produced, we recomputed orthogroup-level evidence summaries from the local files. BLAST and Pfam results were summarized by number of query sequences with hits, top hit labels, domain labels, and best E-values. SignalP outputs were summarized as counts and fractions of proteins with predicted signal peptides. DeepTMHMM outputs were summarized as globular, secreted, transmembrane, or secreted-plus-transmembrane categories and by transmembrane-region counts. These summaries were written to `recomputed_evidence.tsv` in the comparison output directory.

Model relevance labels used different vocabularies, so each call was mapped to a common ordinal tier: low/background/unlikely, watchlist/uncertain, moderate/possible/plausible, and high/direct. We then calculated per-orthogroup mean score, score range, score standard deviation, overcalling runs, and undercalling runs in `comparison_merged_annotation/model_run_comparison.tsv`. Within-method consistency was measured as exact agreement between normalized tier labels for each pair of repeated runs.

### 4.3 Final annotation merge

The final table was curated from three information sources: the recomputed evidence table, the nine model outputs, and manual biological review of disagreement cases. We favored family-wide domain and topology support over isolated hits. We classified an orthogroup as high relevance only when the evidence was both strong and mechanistically close to calcification chemistry, matrix biology, or surface adhesion. Moderate relevance required plausible but incomplete or indirect evidence. Watchlist status was used for orphan secreted or membrane families and weakly connected signaling or vesicle candidates. Low relevance was assigned to housekeeping, intracellular, unsupported, or weakly annotated families without a specific calcification link. The final output was written to `final_orthogroup_annotations.tsv` in the comparison output directory.

## 5 Conclusion

In this controlled annotation task, agentic LLMs were effective at gathering and presenting evidence across dozens of orthogroups, but their biological conclusions were not stable enough to accept without curation. The same prompt and the same evidence produced runs that varied between one and twelve high-confidence calls, and only three orthogroups survived the conservative merge as direct calcification candidates: an arylsulfatase family (OG0017138), a heparan-sulfate-like sulfotransferase family (OG0020703), and an FG-GAP/integrin-like surface-receptor family (OG0018986). Scientific skills and coding-agent interfaces improved reproducibility and provenance, especially by enabling local evidence aggregation, but they did not remove the need for explicit scoring rules and expert review. The strongest practical design is therefore a hybrid one: deterministic bioinformatics evidence generation, replicated agent interpretation, automated disagreement detection, and conservative expert adjudication before final labels are used for downstream biology.

## 6 Data and Code Availability

All benchmark inputs and outputs analyzed in this study are available at github.com/xiaoyu12/agentic_function_annotations.

## 7 Acknowledgments

The author thanks the developers and maintainers of the bioinformatics tools and databases used as evidence sources, including UniProt, BLAST, Pfam, HMMER, SignalP, and DeepTMHMM.

## 9 Author Contributions

X.Z. designed the benchmark, generated or collected the agent outputs, performed the comparative audit, curated the final annotation table, and wrote the manuscript.

## References

[1] The UniProt Consortium. UniProt: the universal protein knowledgebase in 2025. Nucleic Acids Research, 53(D1):D609–D617, 2025. doi: 10.1093/nar/gkae1010.

[2] Stephen F. Altschul, Warren Gish, Webb Miller, Eugene W. Myers, and David J. Lipman. Basic local alignment search tool. Journal of Molecular Biology, 215(3):403–410, 1990. doi: 10.1016/S0022-2836(05)80360-2.

[3] Jaina Mistry, Sara Chuguransky, Lowri Williams, Matloob Qureshi, Gustavo A. Salazar, Erik L. L. Sonnhammer, Silvio C. E. Tosatto, Lisanna Paladin, Shriya Raj, Lorna J. Richardson, Robert D. Finn, and Alex Bateman. Pfam: The protein families database in 2021. Nucleic Acids Research, 49(D1):D412–D419, 2021. doi: 10.1093/nar/gkaa913.

[4] Sean R. Eddy. Accelerated profile HMM searches. PLoS Computational Biology, 7(10): e1002195, 2011. doi: 10.1371/journal.pcbi.1002195.

[5] Felix Teufel, José Juan Almagro Armenteros, Alexander Rosenberg Johansen, Magnús Halldór Gíslason, Silas Irby Pihl, Konstantinos D. Tsirigos, Ole Winther, Søren Brunak, Gunnar von Heijne, and Henrik Nielsen. SignalP 6.0 predicts all five types of signal peptides using protein language models. Nature Biotechnology, 40:1023–1025, 2022. doi: 10.1038/s41587-021-01156-3.

[6] Jeppe Hallgren, Konstantinos D. Tsirigos, Mads Damgaard Pedersen, José Juan Almagro Armenteros, Paolo Marcatili, Henrik Nielsen, Anders Krogh, and Ole Winther. DeepTMHMM predicts alpha and beta transmembrane proteins using deep neural networks. bioRxiv, 2022. doi: 10.1101/2022.04.08.487609.

[7] Predrag Radivojac, Wyatt T. Clark, Tal Ronnen Oron, et al. A large-scale evaluation of computational protein function prediction. Nature Methods, 10:221–227, 2013. doi: 10.1038/nmeth.2340.

[8] Colin Brownlee, Glen L. Wheeler, and Alison R. Taylor. Coccolithophore biomineralization: New questions, new answers. Seminars in Cell & Developmental Biology, 46:11–16, 2015. doi: 10.1016/j.semcdb.2015.10.027.

[9] Alastair Skeffington, Axel Fischer, Sanja Sviben, Magdalena Brzezinka, Michal Górka, Luca Bertinetti, Christian Woehle, Bruno Huettel, Alexander Graf, and André Scheffel. A joint proteomic and genomic investigation provides insights into the mechanism of calcification in coccolithophores. Nature Communications, 14:3749, 2023. doi: 10.1038/s41467-023-39336-1.

[10] Keisuke Kayano, Kazuko Saruwatari, Toshihiro Kogure, and Yoshihiro Shiraiwa. Effect of coccolith polysaccharides isolated from the coccolithophorid, Emiliania huxleyi, on calcite crystal formation in in vitro CaCO3 crystallization. Marine Biotechnology, 13(1):83–92, 2011. doi: 10.1007/s10126-010-9272-4.

[11] Craig J. Dedman, Nishant Chauhan, Alba González-Lanchas, Chloë Baldreki, Adam A. Dowle, Tony R. Larson, Renee B. Y. Lee, and Rosalind E. M. Rickaby. Exploring proteins within the coccolith matrix. Scientific Reports, 14:31821, 2024. doi: 10.1038/s41598-024-83052-9.

[12] Anthropic. Claude code overview. https://docs.anthropic.com/en/docs/claude-code/overview, 2026. Accessed 18 April 2026.

[13] OpenAI. Codex web. https://platform.openai.com/docs/codex/overview, 2026. Accessed 18 April 2026.

[14] K-Dense AI. Scientific agent skills. https://github.com/K-Dense-AI/claude-scientific-skills, 2026. Accessed 18 April 2026.

[15] Matthew W. Vetting, Subray S. Hegde, Joel E. Fajardo, Andras Fiser, Steven L. Roderick, Howard E. Takiff, and John S. Blanchard. Pentapeptide repeat proteins. Biochemistry, 45(1): 1–10, 2006. doi: 10.1021/bi052130w.

[16] Alexandra M. Schnoes, Shoshana D. Brown, Igor Dodevski, and Patricia C. Babbitt. Annotation error in public databases: Misannotation of molecular function in enzyme superfamilies. PLoS Computational Biology, 5(12):e1000605, 2009. doi: 10.1371/journal.pcbi.1000605.

[17] Anthropic. Introducing claude opus 4.7. https://www.anthropic.com/news/claude-opus-4-7, 2026. Accessed 18 April 2026.

