## Supplemental Tables for "Benchmarking Agentic Large Language Models for Complex Protein-Set Functional Annotation"

### Supplementary Information

Supplementary Table S1 provides the complete final merged annotation table for all 73 orthogroups. Supplementary Table S2 gives the full method-level run summary, and Supplementary Table S3 lists the pairwise run-to-run agreement values underlying Figure 3.

Table S1: Complete final merged annotations for the 73 orthogroups. The table is derived from `final_orthogroup_annotations.tsv`.

| Orthogroup | n | Final function | Relevance | Curated rationale |
|---|---|---|---|---|
| OG0000049 | 502 | Heterogeneous FKBP15/coiled-coil-like expanded family | Low | Very low BLAST/PFAM coverage and mostly globular; no calcification-specific evidence. |
| OG0001332 | 91 | Hint-domain or Hedgehog-Warthog-like membrane autoprocessing proteins | Watchlist | Strong Hint-domain and membrane topology support cell-surface signalling, but not a direct mineral-matrix function. |
| OG0001976 | 73 | Pentapeptide-repeat protein family | Low | Strong pentapeptide-repeat support, mostly non-secreted; no direct calcification evidence in the provided data. |
| OG0002887 | 59 | PIF1-family DNA helicase | Low | Housekeeping genome-maintenance helicase. |
| OG0003717 | 51 | Uncharacterized family | Low | No BLAST/PFAM annotation and weak TM signal only. |
| OG0009246 | 29 | Secreted adhesive or ECM-like protein family with weak collagen/adhesive BLAST support | Moderate | SignalP enrichment and adhesive/collagen BLAST hits make this a matrix candidate, but PFAM support is poor and homology coverage is low. |
| OG0009301 | 29 | Secreted or membrane-anchored proline-rich/SHOCT-like surface protein | Moderate | Very strong SignalP and membrane-anchor topology; RPB1 BLAST is likely low-complexity noise. Good surface/matrix candidate but no specific calcification domain. |
| OG0009816 | 28 | Golgi alpha-1,3-mannosyltransferase-like enzyme | Moderate | Mannosyltransferase PFAM and BLAST support glycosylation of secretory or matrix molecules, an indirect calcification-relevant process. |
| OG0010177 | 28 | Ankyrin-repeat scaffold protein | Low | Intracellular scaffold-like annotation with no secretion or calcification-specific evidence. |
| OG0010867 | 27 | Pentapeptide-repeat protein family | Low | Pentapeptide-repeat support but no strong secretion or mineralization-specific evidence. |
| OG0010991 | 27 | Uncharacterized family with minor secretion signal | Watchlist | No functional annotation; 4/27 SignalP positives make it a weak orphan follow-up, not a functional calcification call. |
| OG0011061 | 27 | Exostosin/GT47-like membrane glycosyltransferase candidate | Moderate | All proteins are membrane-associated and two have GT47 support; plausible polysaccharide/matrix biosynthesis role, but low PFAM coverage prevents a high-confidence direct call. |
| OG0011197 | 27 | 2OG-Fe(II) oxygenase or RAP-like protein | Low | Weak oxygenase support and top hits do not indicate matrix hydroxylation or calcification. |

Continued on next page

Table S1: Complete final merged annotations for the 73 orthogroups (continued).

| Orthogroup | n | Final function | Relevance | Curated rationale |
|---|---|---|---|---|
| OG0013037 | 24 | Weak serine hydrolase or aminopeptidase-like family | Low | Sparse PFAM/BLAST support and mixed topology; no calcification-specific signal. |
| OG0014155 | 22 | Secreted or lumenal inosine-uridine nucleoside hydrolase | Watchlist | Strong secretion signal is notable, but enzyme chemistry is nucleotide salvage rather than matrix formation. |
| OG0014250 | 22 | Amidase or Glu-tRNA(Gln) amidotransferase-like enzyme | Low | High SignalP fraction but annotation points to general amidase/translation-related chemistry, not calcification. |
| OG0015153 | 20 | Unresolved multi-pass membrane protein family | Watchlist | Strong membrane topology with poor homology support makes it an orphan membrane candidate. |
| OG0016203 | 18 | Mostly membrane protein family with one collagen/RCC1-like member | Watchlist | One sequence has collagen-repeat evidence and most are TM, but family-wide collagen support is weak. |
| OG0016211 | 18 | SUR7/PalI-family multi-pass membrane organizer | Watchlist | All proteins are TM; membrane organization could matter for vesicle biology but no direct calcification marker is present. |
| OG0017138 | 16 | Arylsulfatase/sulfatase family | High | Strong family-wide Sulfatase PFAM and arylsulfatase BLAST support; sulfated polysaccharide remodeling is directly relevant to coccolith/matrix chemistry. |
| OG0017238 | 16 | tRNA ligase | Low | Housekeeping RNA-processing enzyme. |
| OG0017305 | 16 | Stf0-like sulfotransferase | Moderate | Sulfotransferase PFAM suggests sulfated matrix/glycan chemistry, but only 3/16 PFAM hits and SignalP support is weak by SignalP. |
| OG0017361 | 16 | DJ-1/PfpI family stress or cysteine hydrolase | Low | Stress/metabolic protein family with no specific mineralization link. |
| OG0017362 | 16 | Red chlorophyll catabolite reductase-like protein | Low | Plastid/chlorophyll-catabolism-like evidence despite partial SignalP. |
| OG0017884 | 15 | GNAT acetyltransferase | Low | General acetyltransferase with no calcification-specific evidence. |
| OG0017903 | 15 | Flavohemoprotein or oxidoreductase | Low | Redox/NO-detoxification-like function; no direct calcification evidence. |
| OG0017965 | 15 | Kelch/beta-propeller protein with possible NANM-like domain | Low | Likely intracellular/regulatory scaffold; no secretion or matrix evidence. |
| OG0018519 | 14 | Uncharacterized conserved multi-pass membrane family | Watchlist | 13/14 predicted TM and no homology; worth orphan membrane follow-up. |
| OG0018986 | 13 | FG-GAP/integrin-alpha-like cell-surface receptor with TM or 7TM features | High | Family-wide FG-GAP repeats plus secretion/TM topology support extracellular adhesion/Ca-binding receptor biology, one of the strongest non-enzymatic candidates. |
| OG0019067 | 13 | LRR/NLR-like receptor protein | Low | Immune/receptor-like LRR evidence, not specific to calcification. |
| OG0019078 | 13 | GST or elongation-factor-gamma-like protein | Low | Translation/redox housekeeping signal. |
| OG0019174 | 13 | Amidase or Glu-tRNA(Gln) amidotransferase-like enzyme | Low | Likely housekeeping amidase despite SignalP in some members. |

Continued on next page

Table S1: Complete final merged annotations for the 73 orthogroups (continued).

| Orthogroup | n | Final function | Relevance | Curated rationale |
|---|---|---|---|---|
| OG0019180 | 13 | Uncharacterized soluble/orphan family | Low | No informative homology, domains, secretion, or TM signal. |
| OG0019790 | 12 | HECT E3 ubiquitin ligase with ankyrin repeats | Low | Protein-turnover scaffold, not calcification-specific. |
| OG0019817 | 12 | Secreted or lumenal FAD/NAD monooxygenase-like protein | Watchlist | SignalP enrichment is notable, but homology points to redox/secondary metabolism rather than matrix formation. |
| OG0019873 | 12 | Aminotransferase class I/II | Low | Generic metabolic enzyme. |
| OG0019887 | 12 | Ribosome-associated GTPase or intron maturase-like protein | Low | Organellar/ribosome-associated housekeeping evidence. |
| OG0019917 | 12 | Cyclic-nucleotide-binding or PKA-regulatory-like protein | Watchlist | cAMP signalling can regulate physiology, but evidence is indirect and not calcification-specific. |
| OG0020622 | 11 | RecD2/UvrD-like DNA helicase | Low | DNA repair helicase. |
| OG0020657 | 11 | Uncharacterized partially secreted family | Watchlist | No homology/domain support, but DeepTMHMM and SignalP suggest secretion in a subset. |
| OG0020696 | 11 | Uncharacterized weak membrane/secreted family | Watchlist | No homology/domain support and only partial topology signal. |
| OG0020700 | 11 | PDZ-domain scaffold protein | Low | Generic scaffold with no secretion or mineralization evidence. |
| OG0020703 | 11 | Heparan-sulfate or sulfoglycan sulfotransferase-like enzyme | High | Sulfotransferase BLAST/PFAM plus secretion support direct relevance to sulfated acidic matrix/polysaccharide chemistry. |
| OG0020706 | 11 | Uncharacterized multi-pass membrane family | Watchlist | No homology/domain support but 5/11 DeepTMHMM TM/SP+TM proteins. |
| OG0020728 | 11 | Uncharacterized soluble/orphan family | Low | No informative evidence. |
| OG0020729 | 11 | Uncharacterized soluble/orphan family | Low | No informative evidence. |
| OG0021347 | 10 | Secreted trypsin-family serine protease | Moderate | Strong trypsin PFAM plus secretion; matrix remodeling is plausible but indirect. |
| OG0021406 | 10 | Short-chain dehydrogenase or sugar-metabolism enzyme | Low | Metabolic enzyme with no calcification-specific evidence. |
| OG0021517 | 10 | F-box protein | Low | Protein-turnover adaptor. |
| OG0021520 | 10 | Mostly uncharacterized soluble family with one beta/TM outlier | Low | Topology support is not family-wide. |
| OG0021523 | 10 | PP2C protein phosphatase | Watchlist | Ca/Mg-dependent phosphatase chemistry is peripheral; no direct matrix/transport evidence. |
| OG0021531 | 10 | DegP/HtrA/PDZ protease-like protein | Low | Quality-control protease-like evidence with no SignalP support and no direct calcification link. |
| OG0022355 | 9 | Secreted/membrane DUF2237 protein family | Watchlist | Strong SP/TM topology and DUF support but no known calcification function. |
| OG0022381 | 9 | NADPH-cytochrome P450 reductase-like redox enzyme | Low | ER/redox metabolism, not calcification-specific. |
| OG0022455 | 9 | Uncharacterized soluble/orphan family | Low | No informative evidence. |

Continued on next page

Table S1: Complete final merged annotations for the 73 orthogroups (continued).

| Orthogroup | n | Final function | Relevance | Curated rationale |
|---|---|---|---|---|
| OG0022473 | 9 | Uncharacterized partially secreted/membrane family | Watchlist | Partial topology signal without function; weak orphan candidate. |
| OG0022474 | 9 | F-box protein | Low | Protein-turnover adaptor. |
| OG0022492 | 9 | Beta-propeller/WD40 kinase-associated scaffold | Low | Regulatory scaffold/kinase hits, no calcification-specific evidence. |
| OG0022500 | 9 | Uncharacterized soluble/orphan family | Low | No informative evidence. |
| OG0022520 | 9 | Uncharacterized soluble/orphan family | Low | No informative evidence. |
| OG0022524 | 9 | Pentapeptide-repeat protein family | Low | Pentapeptide-repeat support but no secretion or direct calcification evidence. |
| OG0022528 | 9 | Uncharacterized soluble/orphan family | Low | No informative evidence. |
| OG0023496 | 8 | Protein kinase or GAK-BMP2K-like kinase | Watchlist | Possible vesicle/signalling role, but no direct mineral chemistry or matrix evidence. |
| OG0023566 | 8 | SEY1/RHD3-like ER-fusion GTPase | Low | Secretory/ER morphology role is broad, not calcification-specific. |
| OG0023594 | 8 | GT8 glycosyltransferase-like enzyme | Moderate | GT8 support is consistent with extracellular polysaccharide biosynthesis, a plausible matrix pathway. |
| OG0023657 | 8 | Uncharacterized soluble/orphan family | Low | No informative evidence. |
| OG0023790 | 8 | Red chlorophyll catabolite reductase-like protein | Low | Plastid/chlorophyll-catabolism-like evidence. |
| OG0024846 | 7 | TOD1/MUCI70-like hexosyltransferase | Moderate | MUCI70/TOD1 glycosyltransferase-like domain suggests matrix/cell-wall polysaccharide modification. |
| OG0024979 | 7 | Uncharacterized soluble/orphan family | Low | No informative evidence. |
| OG0025015 | 7 | Uncharacterized pan-calcifier multi-pass membrane family | Watchlist | 7/7 predicted TM and no homology; strong orphan transporter/membrane follow-up candidate. |
| OG0025016 | 7 | Uncharacterized soluble/orphan family | Low | No informative evidence. |
| OG0025059 | 7 | Uncharacterized secreted proline/cysteine-rich orphan protein | Moderate | 5/7 SignalP positives, no TM, no homology; classic novel secreted matrix-candidate profile but no functional domain. |
| OG0026865 | 6 | JmjC/cupin hydroxylase-like enzyme | Watchlist | Hydroxylase chemistry could affect matrix proteins, but evidence points broadly to JmjC/PSR-like enzymes and is indirect. |

Table S2: Full method-level consistency summary. Label distributions were normalized from each agent output.

| Run | Normalized label distribution | Mean score | Score SD | Missing OGs |
|---|---|---|---|---|
| Claude_app_run1 | high=11; low=27; moderate/possible=15; watchlist/uncertain=20 | 1.71 | 1.22 | 0 |
| Claude_app_run2 | high=1; low=67; moderate/possible=5 | 0.68 | 0.64 | 0 |
| Claude_app_run3 | high=8; low=45; moderate/possible=10; watchlist/uncertain=10 | 1.29 | 1.19 | 0 |
| Claude_code_run1 | high=1; low=62; moderate/possible=10 | 0.82 | 0.79 | 0 |
| Claude_code_run2 | high=3; low=33; moderate/possible=15; watchlist/uncertain=22 | 1.36 | 0.95 | 0 |
| Claude_code_run3 | high=2; low=21; moderate/possible=16; watchlist/uncertain=34 | 1.5 | 0.83 | 0 |
| Codex_app_run1 | high=1; low=62; moderate/possible=10 | 0.82 | 0.79 | 0 |
| Codex_app_run2 | low=39; moderate/possible=34 | 1.43 | 1 | 0 |
| Codex_app_run3 | high=12; low=27; moderate/possible=34 | 2.01 | 1.28 | 0 |

Table S3: Pairwise within-method exact agreement of normalized relevance labels.

| Method | Run pair | Exact agreement |
|---|---|---|
| Claude App | R1-R2 | 0.397 |
| Claude App | R1-R3 | 0.603 |
| Claude App | R2-R3 | 0.685 |
| Claude Code + skills | R1-R2 | 0.466 |
| Claude Code + skills | R1-R3 | 0.342 |
| Claude Code + skills | R2-R3 | 0.740 |
| Codex App + skills | R1-R2 | 0.630 |
| Codex App + skills | R1-R3 | 0.411 |
| Codex App + skills | R2-R3 | 0.575 |

Table S4: Representative overclaim patterns identified during manual audit.

| Pattern | Example orthogroups | Conservative interpretation |
|---|---|---|
| Pentapeptide-repeat overinterpretation | OG0001976; OG0010867; OG0022524 | The local evidence supports pentapeptide repeats, not a direct Ca/carbonate-binding or biomineralization function; recent coccolith matrix proteomics nonetheless place pentapeptide-repeat motifs inside the matrix [11], so these orthogroups are reasonable watchlist candidates but not confirmed positives. |
| Secretion treated as sufficient evidence | OG0014250; OG0021347; OG0025059 | Signal peptides nominate extracellular candidates, but function labels must distinguish secreted housekeeping enzymes from matrix proteins and orphan candidates. |
| Single-hit collagen or adhesive transfer | OG0009246; OG0016203 | Matrix-like BLAST hits are useful leads only when supported across the orthogroup or by compatible domains and topology. |
| Low-complexity or giant-protein BLAST labels | OG0009301; OG0011061; OG0015153 | Repetitive BLAST labels such as RPB1 or Piccolo-like hits require downgrading unless matched by domain architecture. |
| Indirect glycan or sulfation chemistry | OG0009816; OG0011061; OG0017305; OG0023594; OG0024846 | Glycosyltransferase and sulfotransferase domains are plausible for matrix chemistry, but partial coverage or weak secretion support should usually remain moderate. |
